# Systematic analysis of the sphingomyelin synthase family in *C. elegans*

**DOI:** 10.1101/2023.07.25.550547

**Authors:** Gaelen G. Guzman, Scotland Farley, Jennifer E. Kyle, Lisa M. Bramer, Sandra Hoeltzl, Joep van den Dikkenberg, Joost C. M. Holthuis, Fikadu G. Tafesse

## Abstract

Sphingomyelin (SM) is a major component of mammalian cell membranes and particularly abundant in the myelin sheath that surrounds nerve fibers. Its production is catalyzed by SM synthases SMS1 and SMS2, which interconvert phosphatidylcholine and ceramide to diacylglycerol and SM in the Golgi and at the plasma membrane, respectively. As the lipids participating in this reaction fulfill both structural and signaling functions, SMS enzymes have considerable potential to influence diverse important cellular processes. The nematode *Caenorhabditis elegans* is an attractive model for studying both animal development and human disease. The organism contains five SMS homologues but none of these have been characterized in any detail. Here, we carried out the first systematic analysis of SMS family members in *C. elegans*. Using heterologous expression systems, genetic ablation, metabolic labeling and lipidome analyses, we show that *C. elegans* harbors at least three distinct SM synthases and one ceramide phosphoethanolamine (CPE) synthase. Moreover, *C. elegans* SMS family members have partially overlapping but also unique subcellular distributions and together occupy all principal compartments of the secretory pathway. Our findings shed light on crucial aspects of sphingolipid metabolism in a valuable animal model and opens avenues for exploring the role of SM and its metabolic intermediates in organismal development.

## Introduction

Sphingolipids are a structurally diverse class of membrane lipids that co-emerged with sterols during the evolution of eukaryotes. They typically represent ∼15% of cellular lipids and are highly enriched in the plasma membrane where they modulate key physical membrane properties, including fluidity, thickness, and curvature(1). Their affinity for cholesterol and ability to self-associate into laterally segregated membrane microdomains that selectively concentrate or exclude membrane proteins to form dynamic platforms that modulate a range of key physiological processes, from phagocytosis and antigen recognition by T cells to synaptic transmission in neurons(2–6). Other sphingolipid species serve as potent signaling molecules; for example, the recently characterized sphingolipid rheostat acts as a central regulator of cell fate: high levels of ceramide (Cer) initiate autophagy and cell death, whereas the conversion of Cer to sphingosine-1-phosphate (S1P) promotes cell survival and is protective against starvation-induced apoptosis(7). However, fundamental questions remain regarding the mechanisms which regulate and mediate the synthesis and metabolism of sphingolipids. The complexity of the sphingolipid metabolic cascade makes it difficult to study this lipid class in higher organism. Thus, understanding the mechanisms of sphingolipid regulation in model organisms such as *Caenorhabditis elegans* (*C. elegans*) may reveal new avenues for unravelling the individual roles that each lipid plays during organismal development and homeostasis.

While the sphingolipid composition varies by cell type, the lion’s share of sphingolipids in mammalian cells accumulate as SM at the exofacial leaflet of the plasma membrane(8). Alongside the major role of SM as a structural membrane lipid, the synthesis and degradation of SM serves as a central mechanism for regulating cellular Cer levels: small changes in the cellular levels of SM can dramatically affect the Cer content of a cell(9, 10). SM production is catalyzed by sphingomyelin synthases (SMS), which transfer the phosphocholine headgroup from phosphatidylcholine to Cer, producing diacylglycerol (DAG) as a side product. DAG is itself a known proliferative signaling molecule(11–13), and thus the production of SM simultaneously regulates Cer and DAG levels in opposing directions – suggesting that SM synthases can play a fundamental role in regulating cell fate towards survival. While numerous reports demonstrate parallel roles for Cer and SM in humans and *Caenorhabditis elegans* (*C. elegans*), to date there have been no analyses on the enzymes responsible for SM synthesis in this model organism(14–17).

We previously reported the identification of a conserved family of polytopic membrane proteins that display the classical characteristics of SMS enzymes in animals capable of SM production(18). Strikingly, this study revealed multiple SMS genes in each of the organisms analyzed, suggesting a powerful selective pressure towards redundancy in SM synthesis. Human cells contain two isoforms of SM synthase, namely SMS1 in the Golgi complex and responsible for producing the bulk of cellular SM, and SMS2 at the plasma membrane and likely serving a principle role in signal transduction(19). In addition, the human genome encodes a third, SMS-related enzyme (SMSr) that catalyses the production of the SM analog ceramide phosphoethanolamine (CPE) in the ER(18, 20, 21). All three enzymes significantly contribute to the local regulation of ceramide levels and serve critical roles in cell growth and survival(19, 20, 22).

The identification of a multigenic SMS family opened important new avenues for studying sphingolipid function in animals. In this regard, *C. elegans* represent an attractive model system. The availability of a detailed description of this animal’s morphology, development and physiology combined with the ease to manipulate gene function through mutation and RNAi makes this model ideal for dissecting the biological roles of SM synthases and related enzymes at the molecular level. In many cases where mammalian systems are too complicated to obtain clear information on the molecular ordering of signaling pathways, *C. elegans* has been instructive. This is particularly true for the mechanisms controlling apoptosis and cell division, two integral and invariant components of *C. elegans* development(23, 24). The *C. elegans* genome encodes five SMS family members, but none of these have been characterized in any detail.

In this study, we carried out the first systematic analysis of SMS family members in *C. elegans*. Combining heterologous expression approaches with combinatorial knockout strategies, we established that several of these homologs function as bona fide SM and CPE synthases with unique subcellular distributions. Our findings lay the foundation for future in-depth investigations of the full regulatory potential of SM synthases and related enzymes in animal growth and development.

## Results

### Phylogenetic analysis and expression of SMS family members in *C. elegans*

The *C. elegans* genome encodes five SMS family members, which we named SMSα, SMSβ, SMSγ, SMSδ, and SMSr. These proteins contain six membrane spans, share a common LPP-like active site motif, and display at least 24% sequence identity with the human SM synthase SMS1. Phylogenetically, ceSMSα, ceSMSβ, ceSMSγ and ceSMSδ form clusters with homologues of other nematodes (i.e., *C. briggsae*, *C. remanei*, and *C. brenneri*) that separate from those containing vertebrate SMS1 and SMS2. In contrast, ceSMSr shows a higher degree of conservation and forms a cluster containing both vertebrate and invertebrate homologues (Figure 1A). To determine which of these SMS family members are actively expressed in *C. elegans*, we performed RT-PCR on total RNA isolated from a mixed-state population of the wildtype strain N2. This allowed the detection of transcripts for ceSMSα, ceSMSβ, ceSMSγ, and ceSMSr (Figure 1B). In contrast, we were unable to detect a transcript for ceSMSδ. The same was true when RT-PCR was performed with an independent SMSδ-specific primer set. This precluded cloning of the ceSMSδ cDNA. For the remainder of this study, we therefore focused on the characterization of ceSMSα, ceSMSβ, ceSMSγ, and ceSMSr.

**Figure 1.**
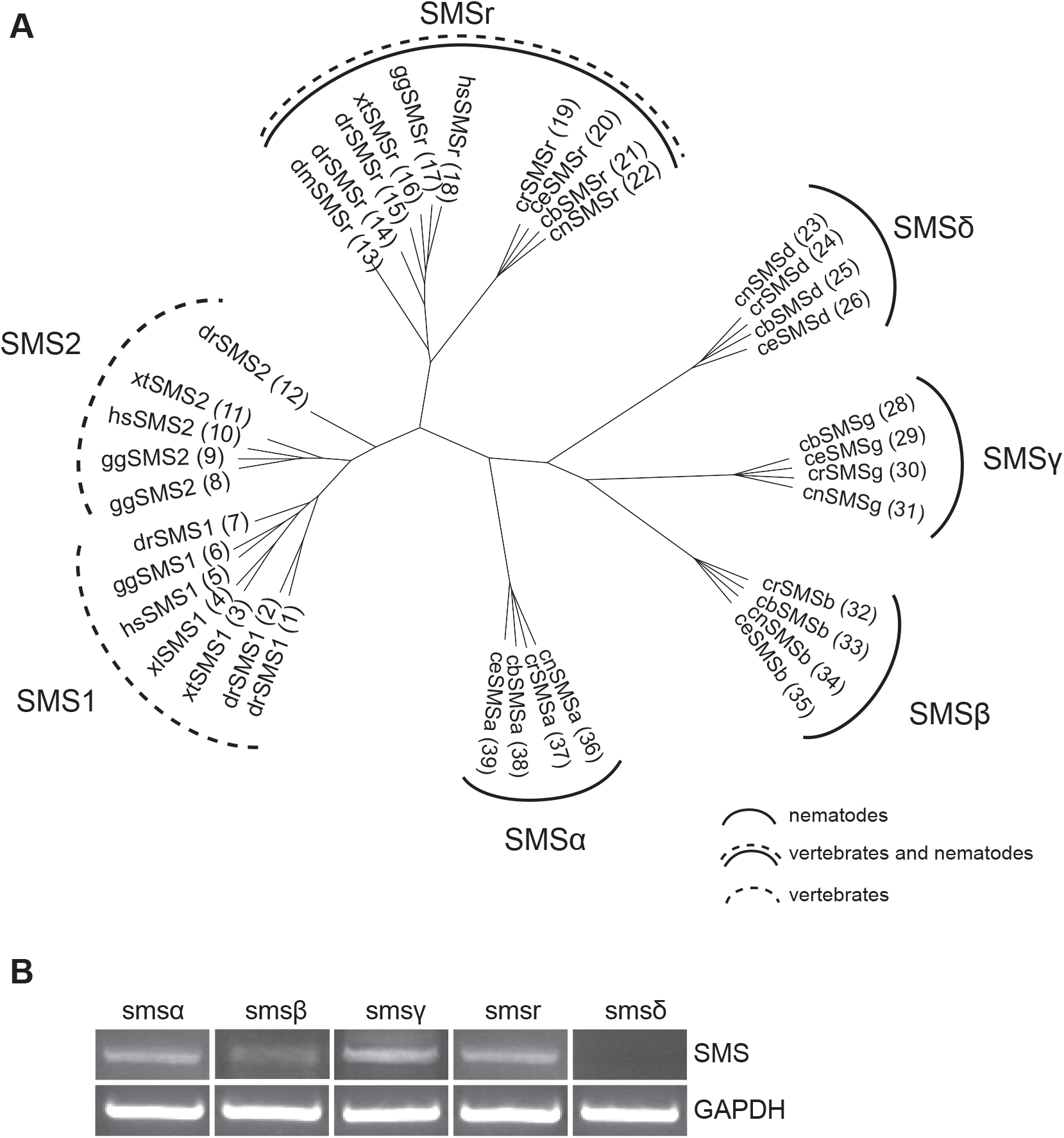
Phylogenetic analysis of SM synthase family and expression levels in *C. elegans*. (**A**) Phylogenetic Tree of SM synthase proteins from *Homo sapiens* (hs), *Drosophila melanogaster* (dm), *Gallus Gallus* (gg), *Xenopus tropicalis* (xt), *Xenopus laevis* (xl), *Danio rerio* (dr) and the nematodes *Caenorhabditis elegans* (ce), *Caenorhabditis briggsae* (cb), *Caenorhabditis remanei* (cr) and *Caenorhabditis brenneri* (cn). (**B**) Expression of SM synthase homologues in *C. elegans* using RT-PCR. Reverse transcription PCR of mRNAs of SMSα, SMSβ, SMSγ, SMSδ and SMSr using total RNA isolated from *C. elegans* N2 worms. To verify that same amounts of RNA were used for each reaction, the *gapdh* gene is shown.

### *C. elegans* SMS family members display distinct subcellular distributions

Previous work revealed striking differences in subcellular distribution among members of the vertebrate SMS family, with SMS1 residing in the Golgi complex, SMS2 in the Golgi and at the plasma membrane, and SMSr in the ER(18, 20). Thus, one potential explanation for the expansion of SMS family members in *C. elegans* could be a further diversification of the cellular sites of SM production. To investigate this possibility, we expressed V5-tagged versions of ceSMSα, ceSMSβ, ceSMSγ and ceSMSr in human HeLa cells and analysed their subcellular distributions by immunofluorescence microscopy. Intriguingly, ceSMSα, ceSMSβ and ceSMSγ each primarily localized to the plasma membrane (Figures 2A, C, and E). In addition, ceSMSα was also found in internal vesicles that were spread throughout the cytoplasm, did not contain the Golgi marker GM130, and likely correspond to compartments of the endosomal/lysosomal system (Figures 2B, C). Besides its association with the plasma membrane, a substantial portion of ceSMSβ co-localized with GM130, indicating that this protein partially resides in the Golgi (Figure 2D, insets). In contrast, ceSMSγ was found almost exclusively at the plasma membrane (Figures 2E, F). CeSMSr, like its human counterpart, resided in the ER, as evidenced by extensive co-localization with the ER marker calnexin (Figure 2H). From this we conclude that members of the *C. elegans* SMS family have partially overlapping but also unique subcellular distributions, together occupying all principal compartments of the secretory pathway (ER, Golgi, plasma membrane, and endosomes).

**Figure 2.**
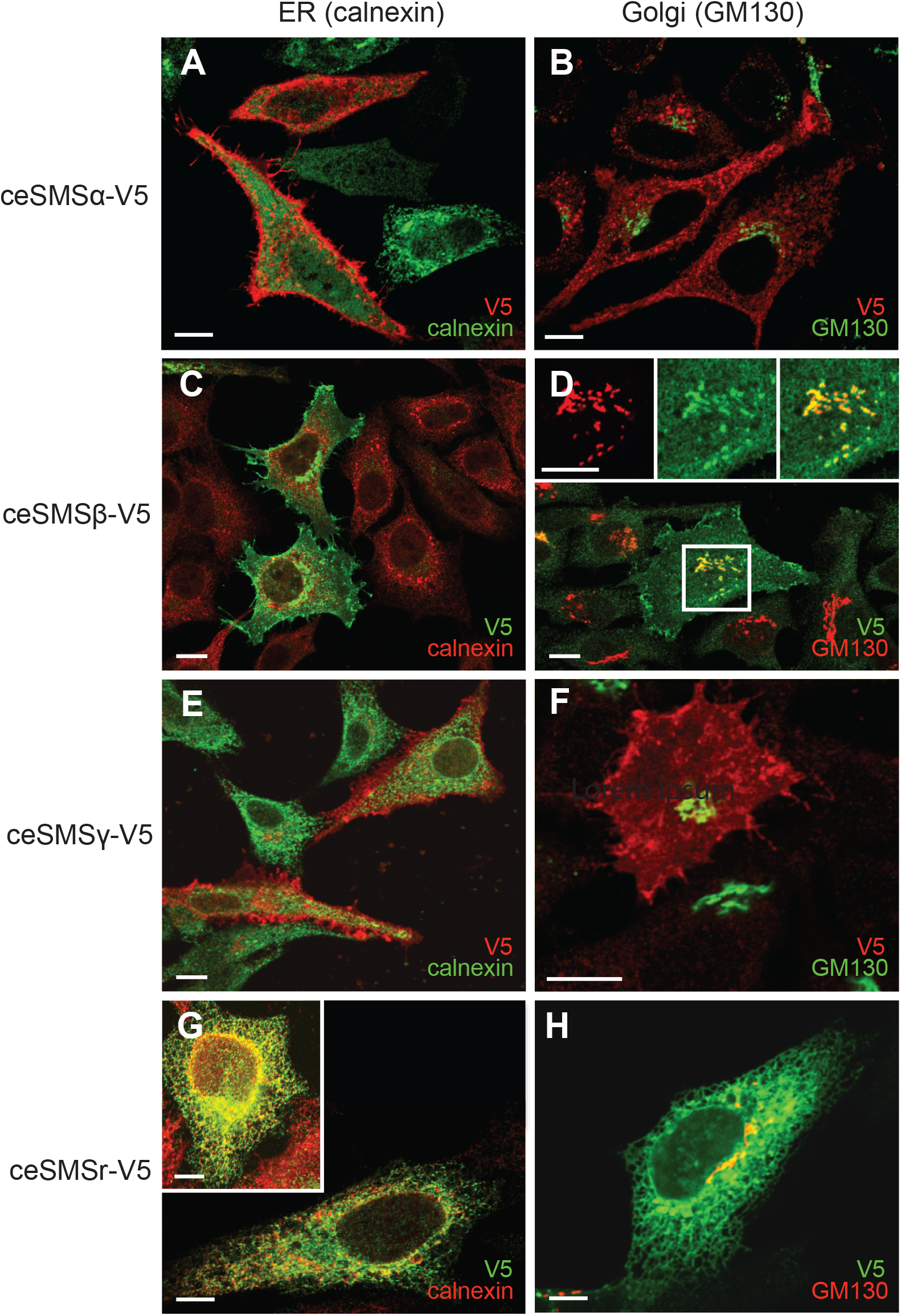
Subcellular localization of *C. elegans* SMS family members in HeLa cells. Confocal immunofluorescence microscopy of HeLa cells transfected with V5-tagged *C. elegans* SMS family members and co-stained with antibodies against the V5 epitope and ER marker calnexin or Golgi marker GM130. Representative images are shown for HeLa cells transfected with ceSMSα-V5 (A, B), ceSMSβ-V5 (C, D), ceSMSγ-V5 (E, F), and ceSMSr-V5 (G, H). Bar, 10 μM.

### The *C. elegans* SMS family harbors both SM and CPE synthases

While human hsSMS1 and hsSMSr primarily serve as monofunctional SM and CPE synthases, respectively, hsSMS2 displays dual activity as SM and CPE synthase(18, 20, 25). To analyze the enzymatic properties of the four *C. elegans* SMS homologues, V5-tagged ceSMSα, ceSMSβ, ceSMSγ and ceSMSr constructs were each expressed in budding yeast, an organism that produces ceramide phosphoinositol (CPI) but lacks endogenous SM and CPE synthase activity. Expression was verified by immunoblot analysis using an antibody against the V5 epitope (Figure 3A). ceSMS-expressing yeast cells were lysed and then incubated with fluorescent C_6_-NBD-ceramide (NBD-Cer). Thin layer chromatographic (TLC) analysis of the reaction mixture derived from ceSMSr-expressing cells showed the presence of an NBD-labeled product with a retention value distinct from that of C_6_-NBD-SM or C_6_-NBD-CPI but similar to that of C_6_-NBD-CPE (Figure 3B). This product was missing in reactions performed with control (empty vector, EV) cells but present in reactions carried out with cells expressing hsSMS2, hsSMSr or *D. melanogaster* drSMSr (Figure 3B and data not shown). These results indicate that *C. elegans* SMSr, like its human and *Drosophila* counterparts, functions as CPE synthase. TLC analysis of the reaction mixtures from cells expressing ceSMSα or ceSMSβ in each case revealed the presence of C_6_-NBD-SM (Figure 3C). Reaction mixtures from cells expressing SMSγ, on the other hand, were devoid of NBD-SM or NBD-CPE. However, all reaction mixtures, including those derived from mock-transfected yeast cells, contained C_6_-NBD-CPI, which is due to the presence of an endogenous CPI synthase. Together, these results indicate that SMSα and SMSβ function as SM synthases.

**Figure 3.**
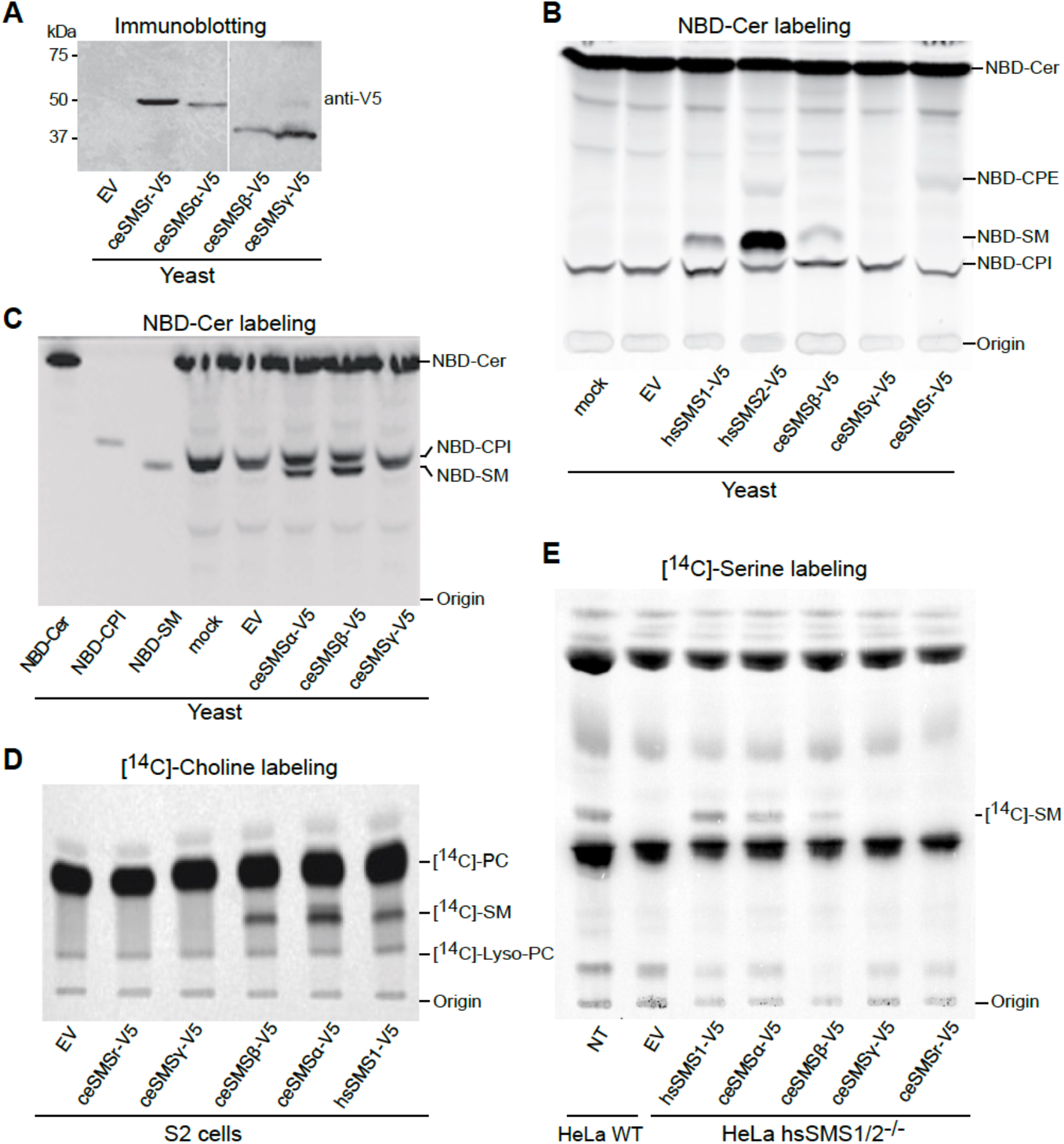
Heterologous expression of C. elegans SMS proteins reveal SM and CPE synthase activity. (**A**) Expression of ceSMS-V5 proteins in yeast cells. Immunoblot of membrane protein extracts derived from yeast cells, transfected with empty vector (EV) or V5-tagged versions of C. elegans *SMS proteins. Constructs were probed with mouse anti-V5 monoclonal antibody. (**B**) SM synthase and (**C**) CPE synthase activity in yeast cells (NBD-Cer labeling). Post-nuclear supernatants of yeast strains expressing human, drosophila and* C. elegans *SMS, or transformed with empty vector (EV) were incubated with NBD-Cer. NBD-labeled lipids were separated by TLC and detected by fluorescence scanning.* (**D**) *Drosophila* S2 cells expressing the different SMS constructs were labeled with [^14^C]-Choline, after which total lipids were extracted and analyzed by radiographic TLC. (**E**) Radiographic TLC showing the metabolic conversion of [^14^C]-Serine in either wild-type or SMS1/SMS2 double-knockout HeLa cells transfected with hsSMS1 or *C. elegans* SMS genes. ‘Origin’ denotes the loading axis; expected retention values for NBD-Cer, NBD-CPE, NBD-CPI, NBD-SM, as well as [^14^C]-PC, [^14^C]-SM, and [^14^C]-lyso-PC, are indicated.

To verify the substrate specificities of *C. elegans* SMS family members and exclude the possibility that the lack of a detectable SM and/or CPE synthase activity of ceSMSγ is due to the use of yeast as a heterologous expression system, we next analyzed the ability of these proteins to support SM production in *Drosophila melanogaster* S2 cells. In comparison to budding yeast, *Drosophila* is evolutionarily closer related to *C. elegans* but also offers the advantage of having no endogenous SM synthase activity. We therefore expressed V5-tagged ceSMSα, ceSMSβ, ceSMSγ and ceSMSr in S2 cells and performed metabolic labeling with [^14^C]-choline to monitor the production of [^14^C]-labeled SM. S2 cells expressing hsSMS1 served as control. TLC analysis and autoradiography of labeled cell extracts showed that hsSMS1, ceSMSα and ceSMSβ each supported SM production (Figure 3D), consistent with the outcome of the experiments in yeast. In contrast, expression of ceSMSγ did not result in any detectable SM production in S2 cells.

Finally, we analyzed the enzymatic properties of *C. elegans* SMS family members in HeLa SMS1/2 double knockout cells that lack endogenous SM synthase activity. Toward this end, the SMS1/2^-/-^ cells were transfected with V5-tagged ceSMSα, ceSMSβ, ceSMSγ and ceSMSr, metabolically labeled with [^14^C]-serine, and then subjected to TLC analysis and autoradiography. SMS1/2^-/-^ cells transfected with hsSMS1 served as control. In line with metabolic labeling experiments in S2 cells, this confirmed that ceSMSα and ceSMSβ function as SM synthases (Figure 3E). In contrast, heterologous expression of either ceSMSγ or ceSMSr did not restore SM production in the SMS1/2^-/-^ cells.

### Lipid profiling of ceSMS-expressing HeLa SMS1/2^-/-^ mutant cells

To interrogate the global impact of individual ceSMS family members on the cellular lipidome, we next performed quantitative lipidomics analysis on HeLa SMS1/2^-/-^ cells transfected with V5-tagged ceSMSα, ceSMSβ, ceSMSγ and ceSMSr. SMS1/2^-/-^ cells transfected with HA-tagged hsSMS1 served as control. Cellular lipids were extracted by the Bligh and Dyer method, spiked with deuterated internal standards, and analyzed with electrospray ionization tandem mass spectrometry (LC-ESI-MS/MS). Lipid extracts from untransfected SMS1/2^-/-^ mutant cells were used to derive ratiometric comparisons against each ceSMS transfected condition and against wildtype HeLa cells.

In our analysis, we identified a total of 537 unique lipid species across all samples, spanning the major lipid classes, namely sphingolipids, phospholipids, glycerolipids, and sterols. The lipid profile of the untransfected SMS1/2^-/-^ cells, as anticipated, exhibited a significant decrease in SM species compared to the wild type HeLa cells (Supplemental Figure 2). Remarkably, upon transfection of SMS1/2^-/-^ cells with hsSMS1, there was about 7-fold increase in total SM levels relative to the non-transfected control (Figure 4A, F, Supplemental Table 1).

**Figure 4.**
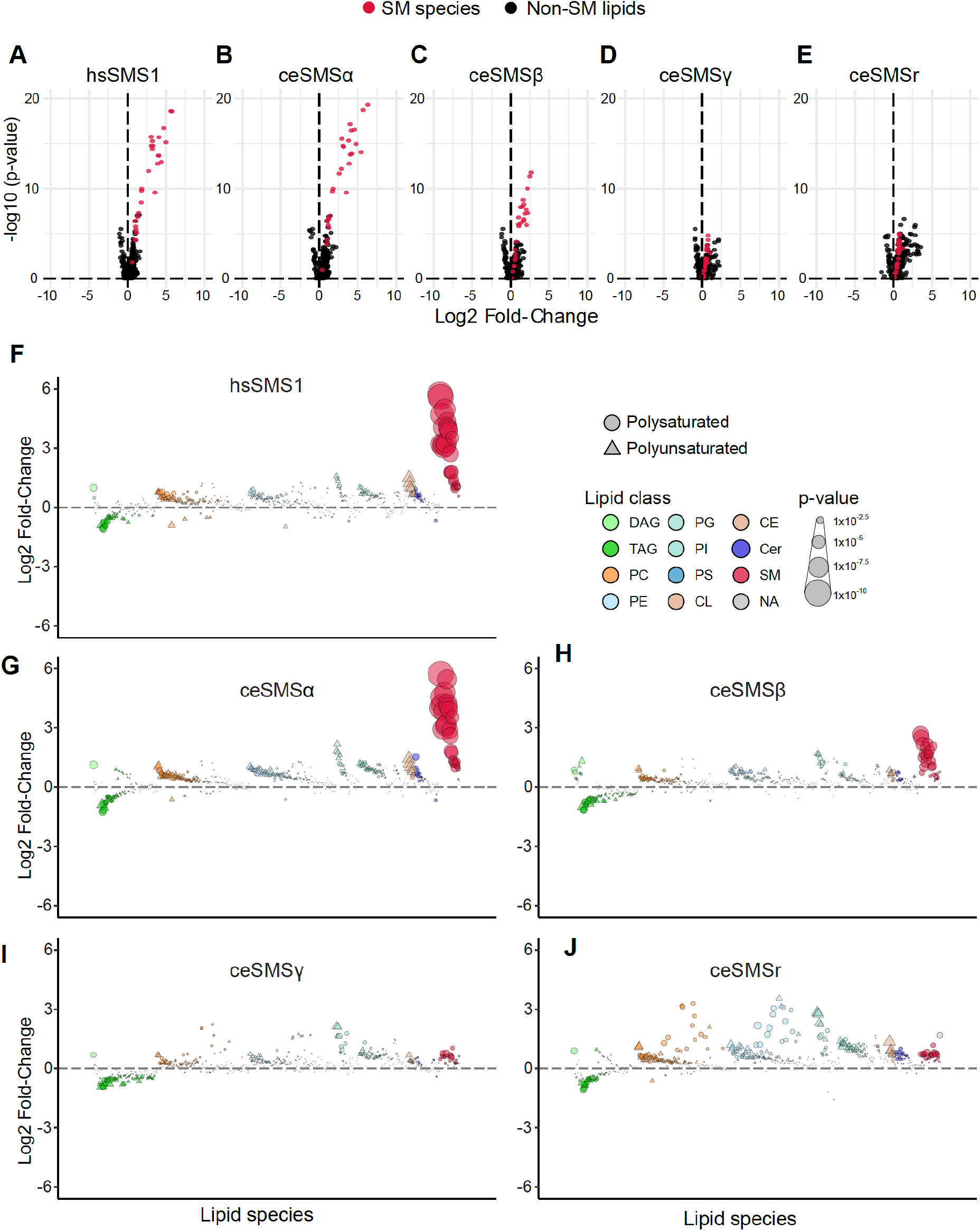
*C. elegans* SMS family members restore SM biosynthesis in SMS1/2-deficient HeLa cells. (**A-E**) Volcano plots depicting the relative abundances and adjusted p-values of lipids quantified via lipidomic analysis, with ratiometric comparisons between transfected and untransfected SMS1/2^-/-^ HeLa cells. Identified SM species are highlighted in red. (**F-J**) Bubble plots of log2 fold changes in abundance of lipid species in SMS1/2^-/-^ HeLa cells transfected with SMS constructs relative to non-transfected controls. Data points are means from five biological replicates; each data point represents a lipid species. Only significantly changed (P < 0.05, one-way ANOVA, with Benjamini–Hochmini adjustment for multiple comparisons) lipids are shown. Individual lipid species are colored by the class of lipid that they belong to. DAG diacylglycerol; TAG triacylglycerol; PC phosphatidylcholine; PE phosphatidylethanolamine; PG phosphatidylglycerol; PI phosphatidylinositol; PS phosphatidylserine; CE cholesterol ester; CL cardiolipin; Cer ceramide; SM sphingomyelin.

Interestingly, when ceSMSα was overexpressed in SMS1/2-deficient cells, the lipid profile pattern closely resembled that of the SMS1/2^-/-^ cells transfected with hsSMS1, and resulted in approximately 8-fold increase of total SM in comparison to the SMS1/2^-/-^ cells (Figure 4B, G, Supplemental Table 1). In a similar manner, the overexpression of ceSMSβ in SMS1/2^-/-^ cells resulted in lipid profiles comparable to those of ceSMSα and hsSMS1 expressing SMS1/2^-/-^ cells, causing an increase of about 2.5-fold in total SM level (Figure 4C, H, Supplemental Table 1). In contrast, overexpressing either ceSMSβ or ceSMSr in SMS1/2^-/-^ cells failed to yield any detectable SM species (Figure 4D, E, I, J). Furthermore, we observed minor alterations in the lipidomes of these SMS-transfected cells, such as slight increases in PC and PE, paired with mild but consistent reductions in TAG (Figure 4F-J).

To gain further insight into the acyl chain preferences of these SM synthases, we conducted a detailed analysis of the nature of the identified SM species. Similar patterns of enrichment were observed for hsSMS1, ceSMSα, and ceSMSβ: SM species, characterized by long-chain and very-long-chain fatty acids (C16:0 and C:20, C:22, and C24:0, respectively), appeared to be preferentially enriched (Figure 5A, B, C).

**Figure 5.**
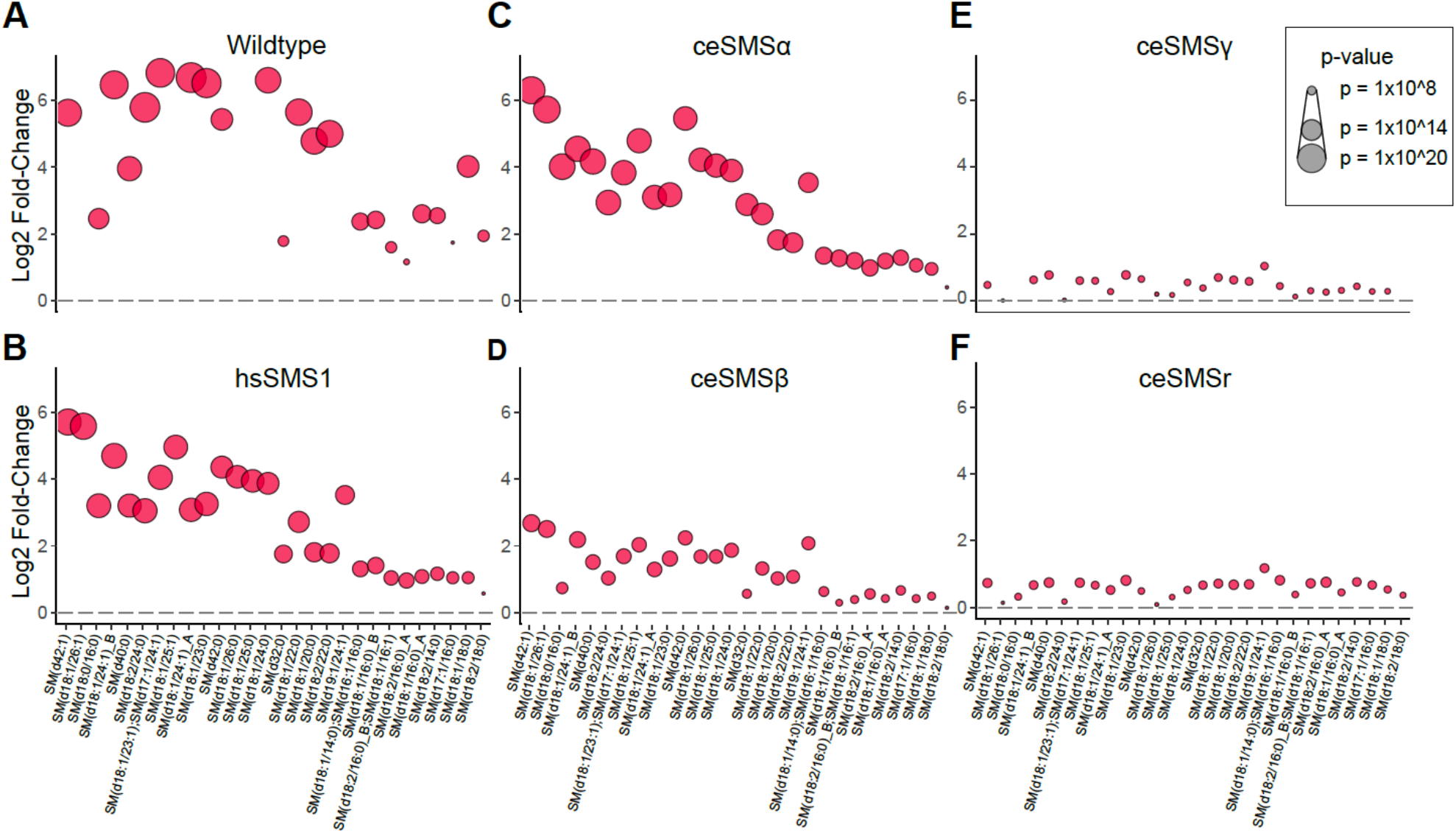
SM species produced by *C. elegans* SMS proteins. Bubble plots of log2 fold changes in abundance of SM species in wildtype HeLa cells (A) and SMS1/2^-/-^ cells transfected with hsSMS (B) or ceSMS constructs (C-F) relative to non-transfected controls. Data points are means from five biological replicates; each data point represents distinct SM species. Only significantly changed (P < 0.05, one-way ANOVA, with Benjamini–Hochmini adjustment for multiple comparisons) lipids are shown. SM species identity is denoted along the X-axis.

Collectively, these data indicate that ceSMSα and ceSMSβ are dedicated SM synthases while ceSMSγ and ceSMSr are devoid of any detectable SM synthase activity. As CPE species largely escaped detection in our lipidomics analysis, further work will be necessary to confirm the CPE synthase activity of ceSMSr detected in our *in vitro* enzyme assays (Figure 3B).

### Characterization of *C. elegans* sms mutants

To analyze the impact of eliminating individual SMS family members in *C. elegans*, deletion mutants in the different *ceSMS* loci were obtained from the National Biosource Project (NBP) in Japan. Mutant strains with deletions in the open reading frames of *ceSMS*α (tm2660), *ceSMS*β (tm2613), *ceSMSδ* (tm2615), and *ceSMSr* (tm2683) genes were obtained. The precise extent and location of the deletions in the *sms* mutants are depicted in Figure 6A and 6B. Mutations were mapped by single-worm PCR on genomic DNA using primers flanking the deletion areas (Figure 6C, upper panel). To check wither the respective deletions in *ceSMS* genes would lead to a complete loss-of-function, the cDNAs corresponding to the mutated *ceSMS* genes were cloned by RT-PCR on total RNA isolated from the *ceSMS* mutant strains (Figure 6C, lower panel).

**Figure 6.**
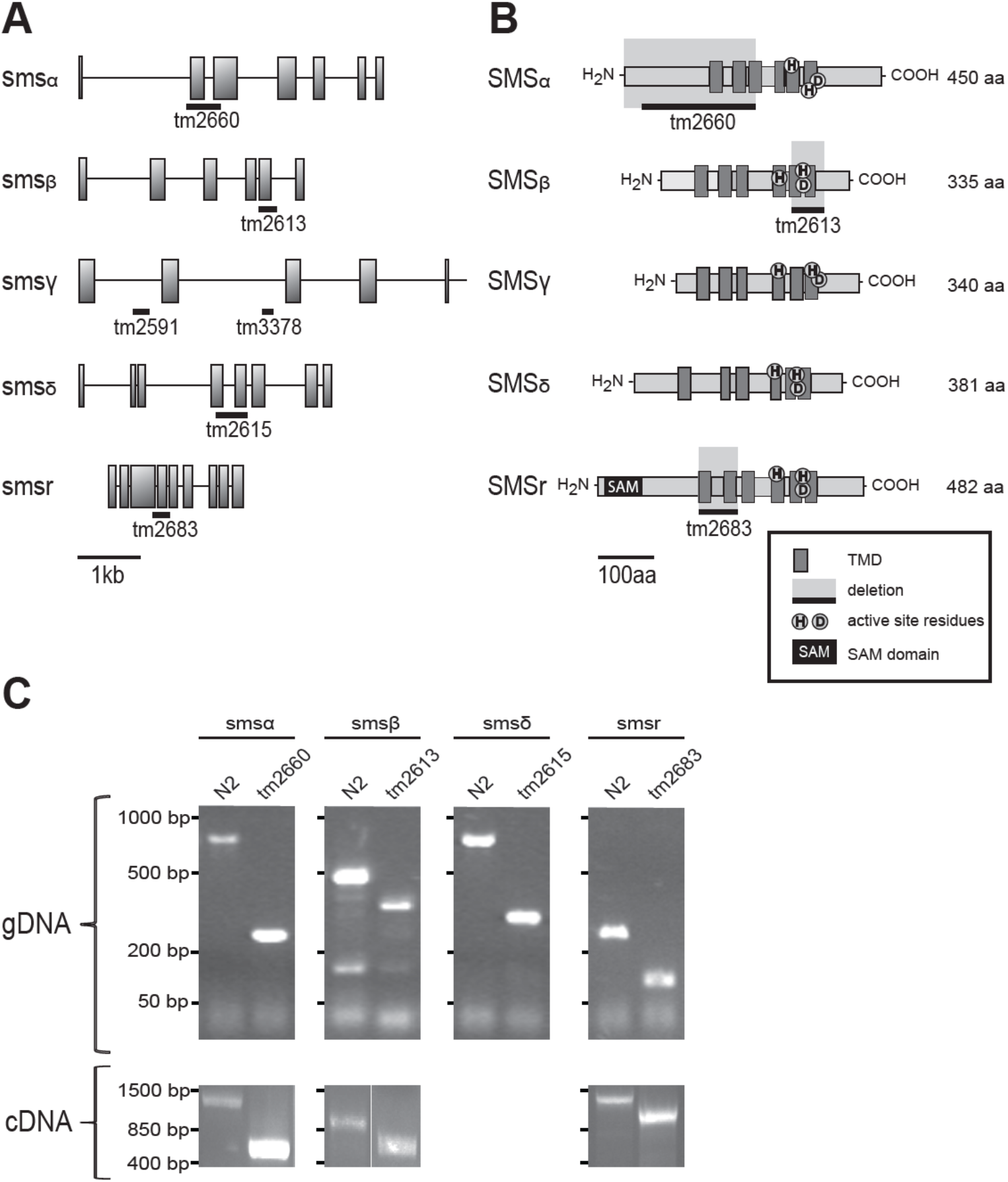
Characterization of SM synthases in *C. elegans*. (**A**) Genetic map of *C. elegans sms* genes. The genomic structures of the *sms-α, sms-β, sms-γ, sms-δ _and sms-r* genes are shown. Gray boxes correspond to coding exons, introns are depicted as lines. The positions of mutations with respect to the gene are indicated below, namely tm2660, tm2613, tm2615 and tm2683. Mutation strains carrying deletions in the *sms-γ* gene, tm2591 and tm3378 have only recently become available, and are not included in this study. (**B**) Schematic structure of the *C. elegans* SMS proteins. The schemes show common domains as the transmembrane domains (solid boxes), the SAM domain (black box) and indicate the position of histidine (H) and aspartate (D) residues, forming the active site of the proteins. The positions of mutations are indicated, and the effect of the deletions on the proteins, are marked by the light gray box. In contrast to the other proteins, the position of the deletion in SMSδ _could not be confirmed by PCR on cDNA. (**C**) Mutant alleles and their mRNA transcripts were characterized by PCR. Locus-specific PCR tests on genomic DNA (gDNA) were used to track deletion alleles. On agarose gels, products from deletion alleles are visibly smaller than the wild type alleles. PCR tests on cDNA reveal a deletion of 588 bp for SMSα, a deletion of 176 bp for SMSβ and a deletion of 210 bp for SMSr. Primers used in this studies are described in Material and Methods.

The tm2660 allele has a deletion of 588 bp that affects two exons in *ceSMS*α (Figure 6A). At the mRNA level, this results in a frameshift at the codon for Gly17 and introduces a premature stop codon after residue 37. The first available start methionine corresponds to Met 232 in wildtype *ceSMS*α. When used, this would give rise to a N-terminally truncated protein that lacks the first three membrane spans (Figure 6B). SMS family members normally have six membrane spanning regions, a feature conserved throughout the LPP superfamily(18). Consequently, it is very unlikely that tm2660 encodes for an enzymatically active version of ceSMSα. A 176-bp deletion in the tm2613 allele leads to a gap in the protein-coding region of *ceSMS*β from residue 234 to 292 (Figures 6A, B). This deletion covers membrane spans 5 and 6 as well as the conserved active site residues His339 and Asp 343. From this, we conclude that tm2613 corresponds to a total loss-of-function allele of ceSMSβ. A 507-bp deletion in tm2615 leads to removal of membrane spans 4 and 5 as well as active site residue His 255 in *ceSMS*δ. This indicates that tm2615 corresponds to a loss-of-function allele of *ceSMS*δ. Finally, a 210-bp deletion in tm2683 causes a gap in the protein-coding region of *ceSMS*r that covers the first two membrane spans. In view of the conserved membrane topology of SMS and LPP proteins, this most likely results in a complete loss of enzymatic activity. Unfortunately, no loss-of-function alleles could be obtained for *ceSMS*γ.

All four of *ceSMS* mutants were homozygous viable, and their overall growth and morphology were indistinguishable from that of wildtype. To eliminate background mutations, each mutant was crossed four times to the wildtype N2 strain. Next, we analyzed the impact of each mutation on SM synthase activity in *C. elegans* using both *in vitro* and *in vivo* assays.

### *C. elegans* contains at least three distinct SM synthases

The foregoing heterologous expression studies in yeast and HeLa cells established that both ceSMSα and ceSMSβ function as SM synthases. To determine the contribution of each enzyme to SM biosynthesis in *C. elegans*, we incubated total worm lysates from corresponding deletion mutants tm2660 (Δsms-α) and tm2613 (ΔceSMSβ) with NBD-Cer and monitored the formation of NBD-SM via TLC. Lysates of the wild-type N2 strain served as control. This showed that disruption of ceSMSα or ceSMSβ in each case resulted in a moderate (20-30%) yet consistent decrease in SM synthase activity (Figure 7A). In contrast, deletion mutants tm2615 (ΔceSMSδ) and tm2683 (ΔceSMSr) showed no reduction in SM synthase activity. Together, these results indicate that both ceSMSα and ceSMSβ contribute to SM synthase activity in *C. elegans*.

**Figure 7.**
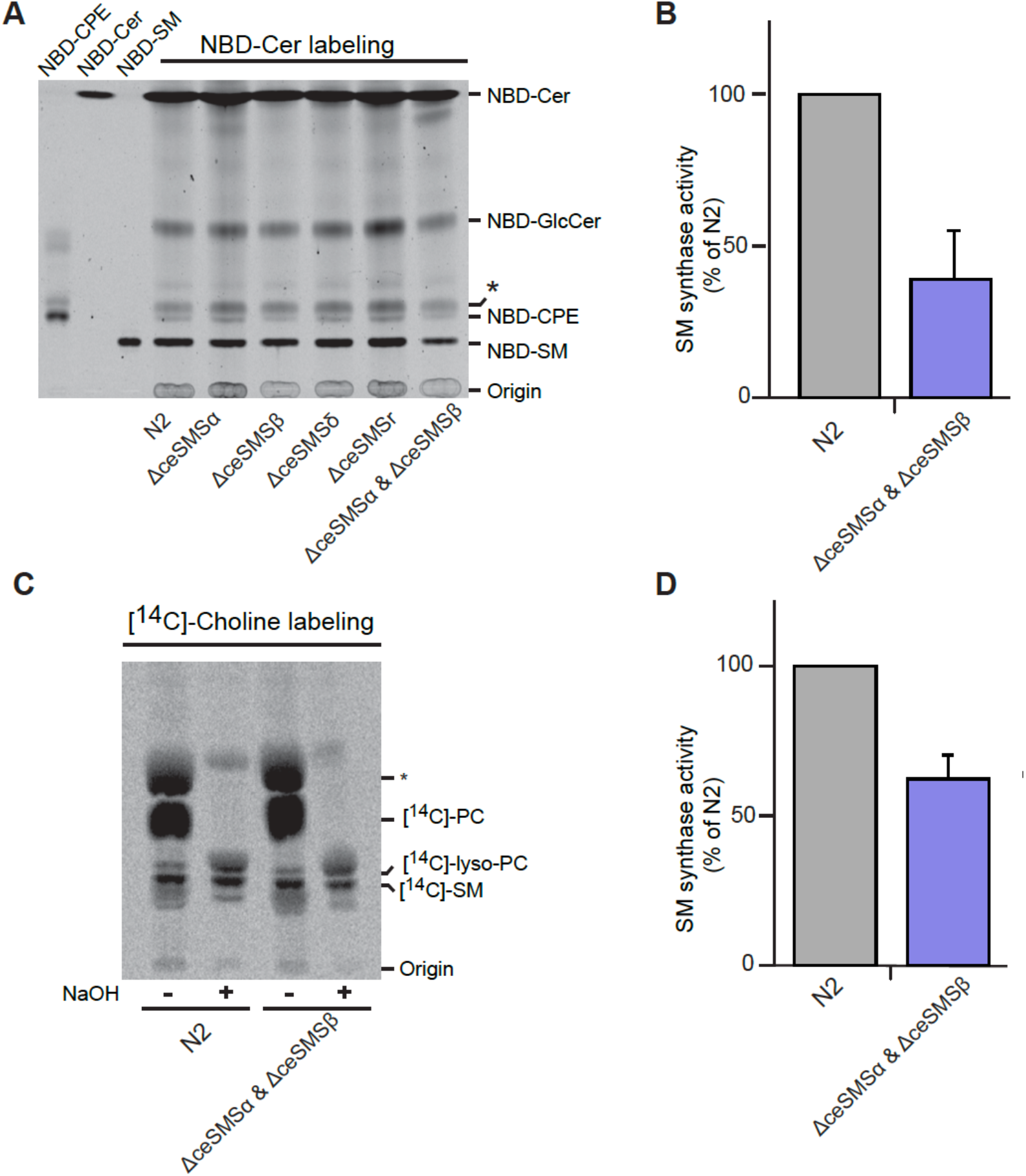
At least 3 different SM synthases contribute to SM synthesis in *C. elegans* worm. (**A**) TLC analysis of reaction products formed, when worm lysates derived from N2 and single or double sms mutant strains were incubated with NBD-Cer for 1 h at 20°C. (**B**) SM synthase activity levels were determined based on quantitative analysis of NBD-SM reaction products formed in worm lysates incubated with NBD-Cer as in A and expressed relative to activity levels in wild type N2 worms. Data are means ± SD, n=5. (**C**) Wild type N2 worms and double mutant strains were labeled with [^14^C]-Choline for 5 h and subjected to lipid extraction, TLC analysis and autoradiography. Glycerolipids were deacylated by mild alkaline hydrolysis (NaOH). (**D**) Quantification of SM synthase activity levels of experiments done as in *D*. Unknown by-products are marked with an asterisk (*). Error bar: range, n=2.

To investigate whether ceSMSα and ceSMSβ represent the only SM synthases in *C. elegans*, we generated a ΔceSMSα ΔceSMSβ double-mutant by crossbreeding tm2660 and 2613. This double mutant exhibited normal growth, was viable, and did not show any noticeable behavioral phenotypes. Interestingly, the double mutant displayed a ∼70% reduction in SM synthase activity compared to the N2 wildtype (Figures 7A, B). To investigate the contribution of ceSMSα and ceSMSβ to SM biosynthesis *in vivo*, the corresponding single and double mutants were metabolically labeled with [^14^C] choline for 4 hr at 20°C and the amount of [^14^C]-labeled SM synthesized was quantified by TLC and autoradiography. Loss of either ceSMSα or ceSMSβ caused only a minor drop (10-30%) in newly-synthesized SM levels, while worms lacking both ceSMSα and ceSMSβ showed a further drop in *de novo* SM synthesis – yet still retained 60% of the wildtype level (Figure 7C, D).

Together, these results provide complementary evidence that ceSMSα and ceSMSβ both operate as SM synthases in *C. elegans*. They also indicate that, besides ceSMSα and ceSMSβ, *C. elegans* harbors at least one additional SM synthase that contributes to SM biosynthesis.

## Discussion

In this study, we present the first systematic analysis of the SMS protein family in the nematode *C. elegans*. The use of heterologous expression systems and gene KO approaches enabled us to assign primary catalytic functions to three of the five SMS family members and map their intracellular sites of action. We show that *C. elegans* ceSMSr catalyzes the production of CPE in the ER, hence analogous to SMSr homologues in mammals and *Drosophila*(*20, 25*). ceSMSα and ceSMSβ, on the other hand, primarily function as SM synthases at the plasma membrane. However, a substantial amount of SMSα resides in endosomes whereas SMSβ partially localized to the Golgi complex. Thus, SMSβ may well represent the *C. elegans* counterpart of SMS1 in mammals. ceSMSγ was found almost exclusively at the plasma membrane but we were unable to assign any enzymatic activity to this protein. The same was true for ceSMSδ, whose expression during the various stages of *C. elegans* development was below the limit of detection. Genetic ablation of both SMSα and SMSβ caused a substantial reduction in, but did not wipe out SM biosynthesis in the worm. This indicates that *C. elegans* contains at least one additional SM synthase, the identity of which remains to be established.

We employed global lipidomics to evaluate how each of these C. elegans enzymes individually influences the lipid profiles, specifically the SM levels, of HeLa cells with SMS1/SMS2 deletion. In line with our expectations, the most notable and significant alterations in SMS1/2 double knockout cells overexpressing hsSMS1, ceSMSα, and ceSMSβ were observed in the SM species. However, overexpression of either ceSMSγ or ceSMSr in SMS1/2-/-cells did not result in detectable levels of SM. This strongly supports our central finding that ceSMSα and ceSMSβ are the primary SM synthases in C. elegans.

Previous work has shown that SMS and CPE synthases serve critical roles in cell growth and survival in mammals. For example, the SMS1-deficient mouse lymphoid cell line WR19L/Fas is growth inhibited when cultured under serum-free conditions – a phenotype that is rescued with ectopic SMS1 cDNA expression(22). Similarly, our previous works showed that depletion of either hsSMS1 or hsSMS2 resulted in reduced growth in human HeLa cervical carcinoma cells and that hsSMSr depletion triggers mitochondria-induced apoptosis in several human cell lines(19, 20). Intriguingly, siRNA knockdown of either hsSMS1 or hsSMS2 in THP-1-derived macrophages *increased* sensitivity to lipopolysaccharide-mediated apoptosis, suggesting that SM production plays a highly nuanced role in determining cell fate(26). Given these previous findings, it is striking that disruption of SMSr or removal of both SMSα and SMSβ in *C. elegans* did not produce recognizable aberration in organismal growth, morphogenesis, or lifespan. It is possible that the corresponding mutants displayed only minor aberrations which escaped our attention, or that these mutants would become defective in growth given adequately stressful conditions. Notably, none of the mutants described in this study displayed complete deficiency in SM or CPE biosynthesis, thus it is feasible that the residual levels of SM and CPE synthase activities are sufficient to sustain growth and survival. Alternatively, the survival of the described *ceSMS* mutants may depend on the activation of pro-mitogenic and/or anti-apoptotic pathways. And finally, we cannot exclude the possibility that nematodes simply do not rely on SMS proteins for vital cell fate decisions in the same ways that mammals do. The creation of novel mutants lacking multiple SMS family members, combined with RNAi approaches, should allow one to address these different scenarios.

Remarkably, nematodes consistently contain more SMS homologues than all other currently known SM and/or CPE-producing organisms (e.g., mammals, insects, etc). While ceSMSα, ceSMSβ and ceSMSr synthesize SM or CPE, this may not necessarily be the case for ceSMSγ and ceSMSδ. A peculiar feature of *C. elegans* is the animal’s ability to esterify its secreted *N*-glycoproteins and glycosphingolipids with phosphorylcholine(27). This property is shared with filarial nematodes in which production of cholinephosphoryl-substituted oligosaccharides (CPOs) serves to avoid the host immune response and contributes to the chronic nature of diseases caused by these parasites(28). Whether cholinephosphoryl-substituted glycoproteins and glycolipids also have intrinsic activities that contribute to nematode development or physiology is unclear. CPO synthesis is catalyzed by a PC:oligosaccharide cholinephosphoryl transferase(27). Because this enzyme is absent in humans, it offers an ideal therapeutic modality for the treatment of filarial infections(28). The identity of the CPO synthase is not known. Since CPO synthesis is mechanistically similar to SM and EPC synthesis, an attractive possibility is that ceSMSγ and/or ceSMSδ functions as a CPO synthase.

## Materials and Methods

### Chemicals and antibodies

NBD-Cer was obtained from Invitrogen (Leek, The Netherlands). NBD-SM, NBD-GlcCer, NBD-PC, POPE (1-palmitoyl-2-oleoyl-sn-glycerol-3-phosphoethanolamine), and POPC (1-palmitoyl-2- oleoyl-sn-glycerol-3-phosphocholine) were obtained from Avanti Polar Lipids, Inc (Alabaster, AL). NBD-CPE was generously provided by P. Devaux (Institut de Biologie Physico-chimique, Paris, France) and other cold lipids were from Matreya Inc. Methyl-[^14^C] choline chloride was obtained from MP Biomedicals (Santa Ana, CA). All other lipids and chemicals were obtained from Sigma-Aldrich (St Louis, MO). The following antibodies were used in this study: rabbit polyclonal anti-human calnexin (Santa Cruz, Santa Cruz, CA), rabbit anti-drosophila calnexin (Abcam Ltd., Cambridge, MA), mouse monoclonal anti-Golgi130 (BD Biomedicals, Alphen aan den Rijn, The Netherlands), mouse/rabbit polyclonal anti-V5 antibodies (Sigma), rabbit polyclonal anti-biotin (Rockland, Gilbertsville, PA) and mouse monoclonal anti-d120kd (EMD). As secondary antibodies we used: Alexa Fluor 488 conjugated goat anti-rabbit (Invitrogen), Alexa Fluor 568 conjugated goat anti-mouse (Invitrogen), HRP-conjugated goat anti-rabbit (Biorad, Veenendaal, The Netherlands) and HRP-conjugated goat anti-mouse (Perbio, Breda, The Netherlands).

### Phylogenetic analysis of SM synthase family

The phylogenetic analysis of SM synthases in nematodes and vertebrates described in Figure 1 was done as described before (38). The proteins sequences were obtained from different sources. UniProt accession numbers, NCBI reference sequences, zebra fish ZFIN or wormbase protein IDs are as follow, organized by SMS family: SMS1: (1) ZDB-GENE-041114-201 (*D. rerio*); (2) Q5U3Z9 (*D. rerio*); (3) Q640R5 (*X. tropicalis*); (4) NP_001018461.1; (5) Q86VZ5 (*H. sapiens*); (6) Q7T3T4 (*G. gallus*); (7) A0JMNO (*D. rerio*). SMS2: (8) F1NLG9 (*G. gallus*); (9) XP_00123149.2 (*G. gallus*); (10) Q8NHU3 (*H. sapiens*) (11) Q5M7L7 (*X. tropicalis*); (12) Q6DEI3 (*D. rerio*). SMSr: (13) Q9VS60 (*D. melanogaster*); (14) A4QNV5 (*D. rerio*); (15) ZDB-GENE-080219-27 (*D. rerio*); (16) Q28CJ3 (*X. tropicalis*); (17) XP_426501.2 (*G. gallus*); (18) Q96LT4 (*H. sapiens*); (19) RP39824 (*C. remanei*); (20) Q20696 (*C. elegans*); (21) CBP26306 (*C. briggsae*); (22) CN16655 (*C. brenneri*). SMSδ: (23) CN19482 (*C. brenneri*); (24) RP33128 (*C. remanei*); (25) CBP15849 (*C. briggsae*); (26) Q9TYV2 (*C. elegans*). SMSγ: (28) CBP32601; (29) Q965Q4 (*C. elegans*); (30) RP06655 (*C. remanei*); (31) CN14615 (*C. brenneri*). SMSβ: (32) RP15984 (*C. remanei*); (33) CBP01983 (*C. briggsae*); (34) CN23351 (*C. brenneri*); (35) Q20735 (*C. elegans*). SMSα: (36) CN26856 (*C. brenneri*); (37) RP41176 (*C. remanei*); (38) CBP43882 (*C. briggsae*); (39) Q9U3D4 (*C. elegans*)

### DNA constructs

hSMS1, ceSMSα, ceSMSβ, ceSMSγ and ceSMSr cDNAs were cloned into yeast expression vector pYES2.1/V5-His-TOPO and mammalian expression vector pcDNA3.1/V5-His-TOPO (Invitrogen) as described previously(18).For expression studies in *Drosophila* S2 cells, hsSMS1, ceSMSα, ceSMSβ, ceSMSγ and ceSMSr cDNAs were PCR amplified and ligated into the copper-inducible pMT/V5-His B vector (Invitrogen).

### Yeast culture

Yeast strain 4Δ.Lass5 (*MAT*a *ade2-101^ochre^ his3-Δ200 leu2-Δ1 lys2-801^amber^ trp1-Δ63 ura3-52 lag1::TRP1 lac1::LEU2 ydc1::natMX ypc1::kanMX4* p413MET25:*Lass5*)(29) was transfected with human and *C. elegans* SMS cDNAs in pYES2.1/V5-His-TOPO and grown in selective synthetic medium containing 2% (wt/vol) galactose and 50 mg/l myo-inositol.

### Cell culture and transfection

HeLa cells were grown in DMEM with 10% FCS (PAA laboratories GmbH, Pasching, Austria). Using Lipofectamine2000 and Lipofectamine3000 (Invitrogen), cells were transiently transfected with SMS-V5/pcDNA3.1 constructs. SMS1/SMS2 double-knockout HeLa cells were kindly provided by the lab of Joost Holthuis and were grown and transfected as wildtype cells. *Drosophila* S2 cells were grown in Schneider’s insect medium with 10% FBS (Cambrex, Rockland, ME) at 27°C in a humidified atmosphere. Insect cells were transfected with SMS/pMT/V5-HisB constructs using Effectene (Qiagen, Hilden, Germany) following the manufacturer’s protocol. Expression of recombinant SMS protein was induced by addition of 1 mM CuSO_4_ for 3 h followed by a 2 h-chase.

### Nematode culture

Worms were maintained on 10 cm nematode growth medium (NGM) agar plates carrying a lawn of *E. coli* OP50 or in liquid culture (S-medium(30)) supplemented with *E. coli* OP50. Culture plates and liquid cultures were maintained at 20°C. Worm strains used in these experiments were: N2 (wild type), Δ*ceSMSα* (tm2660), Δ*ceSMSβ* (tm2613), Δ*ceSMSr* (tm2683) and Δ*ceSMSδ* (tm2615). All strains were provided by the National Biosource Project in Japan (NBP). All mutant strains have been backcrossed 4 times to N2. Mutant alleles were followed by PCR on single worm lysates, detecting the presence of genomic deletions (588 bp in the case of *tm2660*, 176 bp in the case of *tm2613*, 507 bp in the case of *tm261*5 and 210 bp in the case of *tm2683*). Primers, flanking the deletions alleles used include: *sms-α* / tm2660; ttggctattcactccaccct / tcgaagtccccatggaatct; *sms-β* / tm2613: tctgcttacattgggcacat / tcattaacttggccagtgcag; *sms-δ* / tm2615: tctagggcgtcggttggct / tgtaatcgttgtggagcatac; *sms-r* / tm2683: cgacaagtcgagagacccg / cgggtctctcgacttgtcg.

### Creation of double mutant (Δ*ceSMS*α Δ*ceSMS*β)

Backcrossed Δ*ceSMSα* (tm2660) worms were heat shocked (30°C for 4 h) at L4 larval stage and 6 young resulting male offspring were transferred to a plate containing one single L4 Δ*ceSMSβ* (tm2613) hermaphrodite. The progeny were singled out onto plates and maintained until maturity (identified by the presence of eggs). Following single worm lysis, a genomic polymerase chain reaction (PCR) was performed to screen and identify double knockout mutants. Homozygous double mutant strains were selected and assessed and confirmed over at least 2 generations.

### RT-PCR

RNA of worms was isolated using NucleoSpin RNA II kit (Macherey-Nagel, Düren, Germany) according to manufacturer’s protocol. The amount and quality were verified by gel electrophoresis and spectrophotometry. Single strand complementary DNA (cDNA) was generated from 1 µg RNA using dT-oligo primers and Superscript II reverse transcriptase (Invitrogen). PCR was performed using Phusion DNA polymerase (Finnzymes, Espoo, Finland) on 5 µl of cDNA using the cloning primers previously described in Huitema et al. (2004) except for sms-*α* where the following primer pair was used 5’-*gcggccgc*cttcgaaagcaggtcgtgcagctcc-3’/5’- *ggtacc*aagccatgaaaatgtcttggaatcatcaa-3’.

### Immunofluorescence microscopy

Cells were fixed in 4% paraformaldehyde/PBS, processed for immunofluorescence as described previously for S2(31) and HeLa cells(20), and mounted in Vectashield medium containing DAPI (Vector Laboratories, Burlingame, CA). Images were captured at room temperature using a confocal microscope (Figure 2: LSM 510 Meta; Carl Zeiss, Inc., Sliedrecht, The Netherlands. Supplemental Data Figure 1: LSM900; Carl Zeiss, Inc., Sliedrecht, The Netherlands) with a 63x 1.40 NA Plan Apo oil objective. The fluorochromes used were DAPI, l_ex_ = 360 nm and l_em_= 460 nm; Alexa Fluor 488, l_ex_ = 488 nm and l_em_= 515 nm; Alexa Fluor 568, l_ex_ = 568 nm and l_em_ = 585 nm. Images were captured using EZ-C1 software (Nikon Instruments Europe, Badhoevedorp, The Netherlands) and further processed using either Photoshop (version 7.0.1; Adobe) and FIJI (version 2.1.0/1.53c; ImageJ).

### Western blotting

S2 *a*nd yeast cells were lysed in RIPA buffer (50mM Tris/HCl, 0.1% (wt/vol) SDS, 0.5% (vol/vol) NP40, 150 mM NaCl, 2 mM EDTA, pH 7.4), supplemented with Complete Protease Inhibitor Cocktail (PIC, Roche, Basle, Switzerland). Nuclear DNA was sheared by passing lysates through a 23G needle. Protein content of lysates was determined by the bicinchoninic acid method (Pierce, Breda, The Netherlands). Equal amounts of protein in 1x SDS sample buffer were separated by SDS-PAGE and subsequently transferred to a nitrocellulose membrane (Schleicher & Schuell, Dassel, Germany). Membranes were blocked in TBS (25 mM Tris/HCl, 137 mM NaCl, 2.7 mM KCl, pH 7.4) containing 0.05% (vol/vol) Tween 20 and 4% (wt/vol) dried skim milk (Fluka, Zwijndrecht, The Netherlands) and incubated overnight with primary antibody of interest. Membranes were washed three times in TBS containing 0.05% (vol/vol) Tween20 and incubated for 1 h with secondary antibody. Proteins were detected using HRP-conjugated secondary antibodies and enhanced chemiluminescence (Amersham, Roosendaal, The Netherlands).

### In vitro enzyme assays

100 ODs of yeast cells was lysed by bead bashing in 10 ml ice-cold reaction buffer (0.3 M sucrose, 15 mM KCl, 5 mM NaCl, 1 mM EDTA, 20 mM Hepes-KOH, pH 7.0) containing freshly added protease inhibitors. 200 µl postnuclear supernatant (PNS; 700 *g* for 10 min at 4°C) were combined with 200 µl reaction buffer containing 0.002% Triton X-100, 40 nmol POPE and/or POPC, and 50 µM NBD-Cer (Avanti Polar Lipids, Inc.), and incubated at 37°C for 2 h. 2×10^5^ worms were homogenized in an eppendorf tube with 40 strokes of a Teflon pestle in 300 µl ice-cold reaction buffer. 200 µl of worm lysate were combined with 200 µl reaction buffer containing 25 µM NBD-Cer (Avanti Polar Lipids, Inc.), and incubated at 20°C for 2 h. Reactions were stopped by adding 1 ml MeOH and 0.5 ml CHCl_3_, and lipids were extracted according to Bligh and Dyer(32). The lower phase was evaporated under N_2_ and the reaction products analyzed by TLC, which was developed in CHCl_3_/acetone/MeOH/acetic acid/H_2_O (50/20/10/10/5 [vol/vol/vol/vol/vol]). Fluorescent lipids were visualized on an image analysis system (STORM 860; GE Healthcare) and quantified with Quantity One software (Bio-Rad Laboratories).

### Metabolic labeling

*S2* cells (2–5 x 10^6^) grown in 0.5 ml complete Schneider’s insect medium, SMS1/SMS2 double-knockout HeLa cells were grown in DMEM, and *C. elegans* worms (2×10^5^) grown in liquid culture (S-medium) – all three of these models were subsequently radiographically labeled with 1 µCi [^14^C] choline for different time points as indicated. Lipids were extracted in CHCl_3_/MeOH/10 mM acetic acid (1/4.4/0.2 [vol/vol/vol]) and processed according to Bligh and Dyer(32). Half of the extract was subjected to mild alkaline hydrolysis. Radiolabeled lipids were analyzed by TLC in CHCl_3_/MeOH/25% NH_4_OH (50/25/6 [vol/vol/vol] detected by exposure to imaging screens (BAS-MS; FujiFilm), scanned on a Personal Molecular Imager (Bio-Rad Laboratories), and quantified using Quantity One software.

### Lipidomic analysis – sample preparation

SMS1/SMS2 double-knockout HeLa cells were seeded in 6cm dishes at a density of 2×10^6^ cells per dish and transiently transfected in quadruplicate 24 hrs later with appropriate SMS constructs using Lipofectamine3000 (Invitrogen). At 48 hrs post-transfection, cells were washed twice in ice cold PBS and lipids were extracted using CHCl_3_:MeOH:H_2_O (2:1:1, v:v:v) according to Bligh and Dyer(32)(1)10% of each sample was isolated and subjected to BCA analysis to quantify and normalize to protein content. Samples were each spiked with 8uL heavy standards (EquiSPLASH Mix; Avanti Polar Lipids), dried *in vacuo*, and stored at -80°C in a 2:1 solution of chloroform:methanol (v:v) until analysis.

### Lipidomic analysis – LC-MS/MS analysis and lipid identification

LC-MS/MS analysis and lipid identification were performed as previously described(33, 34). LC-MS/MS instrumentation consisted of a Waters Acuity UPLC H class system interfaced with a Velos-ETD Orbitrap mass spectrometer. Samples were again dried *in vacuo* and resuspended in a 1:54 solution of chloroform:methanol (v:v) and injected onto a reverse-phase Waters CSH column and separated over a 34 min gradient (mobile phase A: acetonitrial/H_2_O (40:60, v:v); mobile phase B: acetonitrile/isopropanol (10:90, v:v) containing 10mM ammonium acetate) at a flow rate of 250 μL/min. Both positive- and negative-mode analyses were employed, using higher-energy collision dissociation and collision-induced dissociation to maximize lipidome coverage. The fragment ions used for lipid identifications were used as previously described(34); in summary, these LC-MS/MS raw data files were analyzed using the LIQUID software(33) and all subsequent identifications were manually validated by examining the fragmentation spectra for diagnostic and fragment ions corresponding to the parent acyl chains. For quantification, a reference database was created using LIQUID(33) and then aligned to a reference database based on their identification, *m/z*, and retention times using MZmine 2(35). Aligned features were then manually verified, and peak apex intensity values were exported for statistical analysis.

In total there were 3 (0.04%) and 272 (3.3%) missing observations in positive and negative mode, respectively; all missing observations were coded as NA values, and all observed abundances were log2 transformed. Lipid abundance profiles for each replicate were visualized via boxplot to identify any samples with substantial differences (Supplemental Figure 2b). We note that a single replicate of ceSMSβ transfection resulted in a lower overall abundance distribution in positive ionization mode. This replicate was the lone replicate flagged as a potential outlier (p-value=0.0001) using a robust Mahalanobis distance algorithm (rMd-PAV) based on lipid abundance vectors(36). This replicate was removed from both ionization modes of data, as only 21 of the 291 lipids were observed in negative mode for this replicate. A total of 9 and 8 heavy internal standards (IS) were spiked into the positive and negative ionization data, respectively. ISs with a coefficient of variation (CV) value less than 20% were used to derive normalization factors. 4 ISs (15:0-18:1(d7) DAG, 15:0-18:1(d7) PC, 18:1(d7) Lyso PE, C15 Ceramide-d7) and 3 ISs (15:0-18:1(d7) PC, 18:1(d7) Lyso PC, C15 Ceramide-d7) were used for the positive and negative data, respectively. The median value of ISs for each sample was calculated and used to normalize the data (Supplemental Figure 2c).

For each lipid, an analysis of variance (ANOVA) model was fit, using the R package pmartR(37), to the data and the following post-hoc pairwise differential expression tests were conducted: ceSMSα vs NT, ceSMSβ vs NT, ceSMSγ vs NT, ceSMSr vs NT, hSMS1 vs NT, and WT vs NT. A Holm multiple test correction(38) was used to adjust p-values for multiple comparisons; these adjusted p-values were then transformed to (-1)*Log10 adjusted p-values for visualization.

## Supporting information

Celegans_LipidomicsResults (.csv)

## Acknowledgements

We thank Philippe Devaux and Andreas Conzelmann for gifts of reagents and cell lines and the National Biosource Project in Japan (NBP) for *C. elegans* strains. We also thank the members of the OHSU Advanced Light Microscopy core for their perennial guidance. J.C.M. Holthuis is supported by grants from the DFG (SFB 1557 – P9). F.G. Tafesse is supported by grants from the National Institutes of Health (NIAID R01 AI141549-02). Lipidomics analyses were performed in the Environmental Molecular Sciences Laboratory, a national scientific user facility sponsored by the Department of Energy (DOE) Office of Biological and Environmental Research located at PNNL. PNNL is a multiprogram national laboratory operated by Battelle for the DOE under Contract DE-AC05-76RLO 1830.

---

**Supplementary Figure 1.**
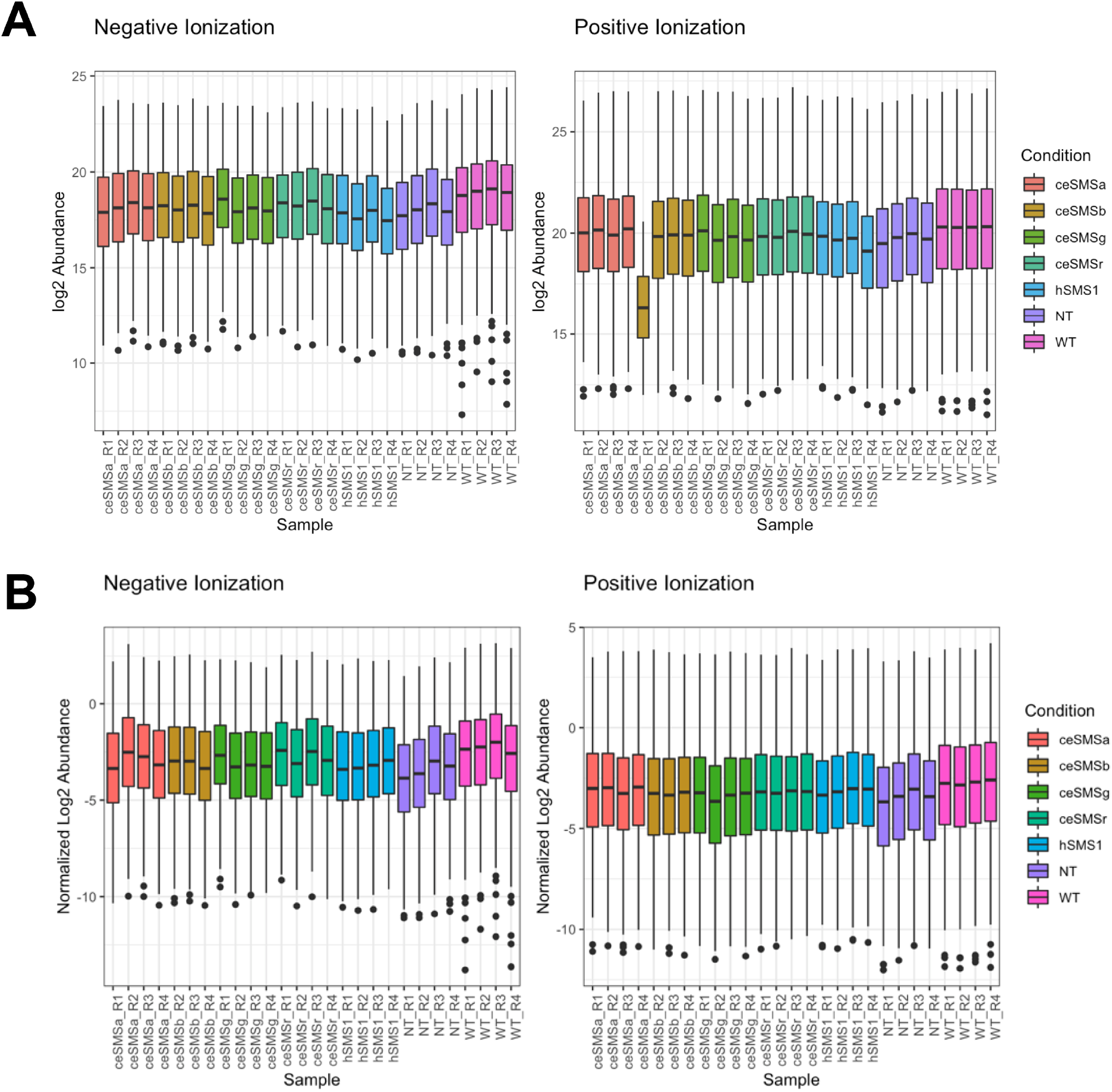
Transfection lipidomics data quality control. (A) Representative confocal microscopy images of SMS1/SMS2 double-knockout HeLa cells transfected with either human SMS1 or ceSMS candidates. Cell bodies were stained with Phalloidin (green), nuclei were stained with DAPI (blue), and transfected enzymes were stained with either anti-HA or anti-V5 (red). (B & C) Boxplots depict distributions of total lipid abundances by replicate before and after normalization via internal standard, highlighting a single replicate of ceSMSβ with an overall lower distribution in positive ion mode.

**Supplemental Figure 2.**
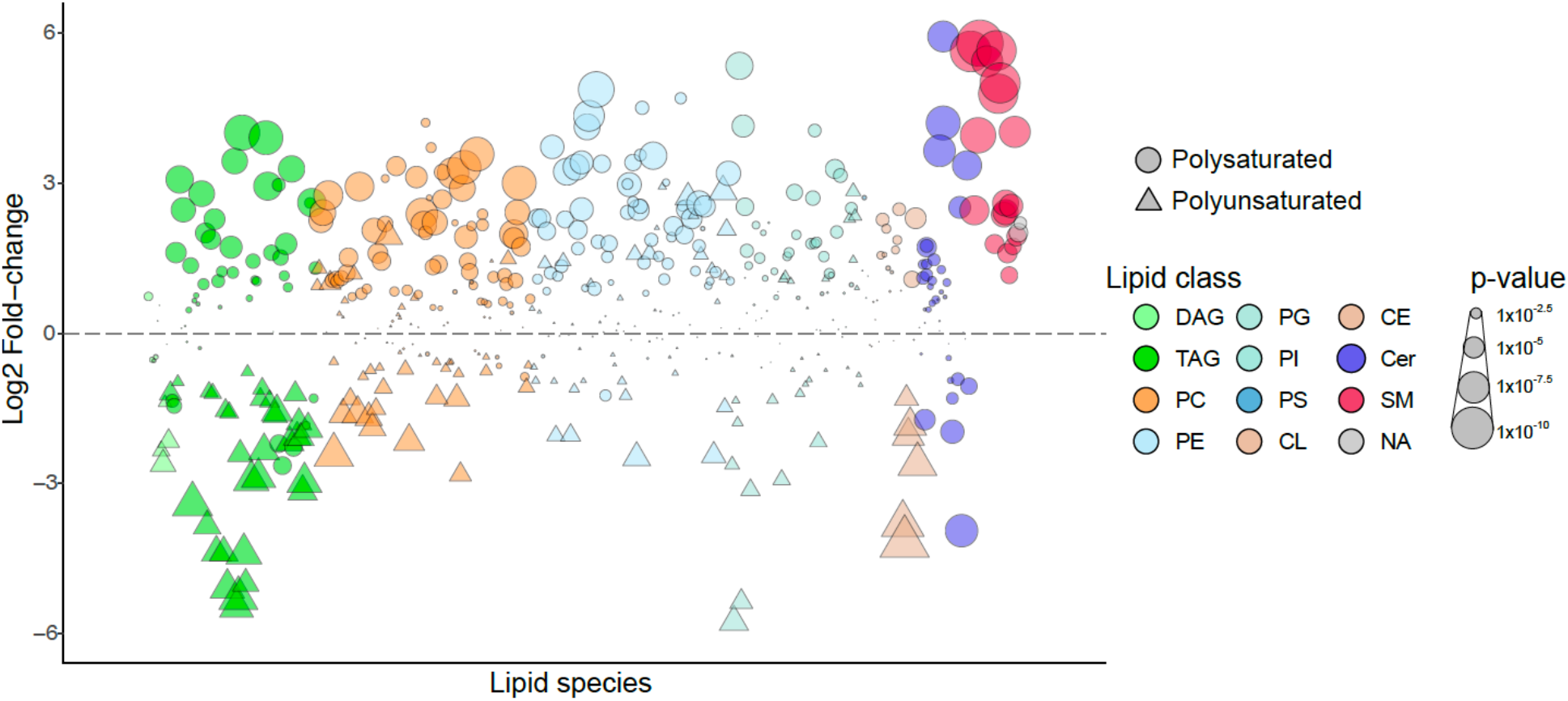
Lipidomics comparison of wildtype and SMS1/2^-/-^ HeLa cells. A bubble plot of log2 fold changes in abundance of lipid species in wildtype HeLa cells relative to SMS1/2^-/-^ cells. Data points are means from five biological replicates. Each individual lipid species is assigned a unique color based on its lipid class. DAG, diacylglycerol; TAG, triacylglycerol; PC, phosphatidylcholine; PE, phosphatidylethanolamine; PG, phosphatidylglycerol; PI, phosphatidylinositol; PS, phosphatidylserine; CL, cardiolipin; Cer, ceramide; HexCer, hexosylceramide; SM, sphingomyelin; CE, cholesterol ester.

## Notes

### Competing Interest Statement

The authors have declared no competing interest.

